# Metabolomic, lipidomic, and N-glycomic analyses of a human cell model of Krabbe disease reveal treatable deficits in glycosylation and serine-ceramide metabolism

**DOI:** 10.64898/2026.08.12.744295

**Authors:** Rodrigo Starosta, Hannah Saeger, Johanna ten Hoeve, SJ Kim, Johan L.K. Van Hove, Xuntian Jiang, Miao He, Neal Bennett

## Abstract

Krabbe disease is a rare autosomal recessive lysosomal disease caused by deficiency of galactocerebrosidase (GALC), leading to accumulation of galactosylceramide and formation of the toxic metabolite galactosylsphingosine (psychosine). While psychosine accumulation is well-established as a primary pathogenic mechanism, the broader metabolic consequences of GALC deficiency remain incompletely understood. In this study, we used stable isotope tracing to comprehensively characterize metabolic perturbations in a human oligodendrocellular Krabbe disease model. This approach revealed elevated *de novo* ceramide synthesis in *GALC* knock-out cells, characterized by increased incorporation of glucose-derived serine into ceramide biosynthetic pathways. This enhanced ceramide production was amenable to pharmacological intervention by tezacaftor, an inhibitor of sphingolipid Δ^4^-desaturate (DEGS); tezacaftor administration also normalized psychosine levels, raising the possibility of its use as substrate reduction therapy. Additionally, we identified significant disruption of UDP-hexose metabolism, manifesting as an overabundance of truncated and hypogalactosylated glycans. These findings suggest impaired protein glycosylation as a previously unrecognized pathogenic mechanism in Krabbe disease. Our findings reveal novel metabolic dysregulation in Krabbe disease extending beyond established psychosine toxicity. The identification of enhanced *de novo* ceramide synthesis presents a new therapeutic target, while the discovery of galactose-deficient glycosylation defects supports galactose supplementation as a potential therapeutic intervention. These metabolic insights provide new mechanistic understanding and therapeutic opportunities for this devastating neurodegenerative disorder.

## INTRODUCTION

Krabbe disease (KD), also known as globoid cell leukodystrophy, is a devastating lysosomal disorder characterized by progressive demyelination of the central and peripheral nervous systems. This autosomal recessive condition affects approximately 1 in 100,000 live births and results from pathogenic variants in *GALC*, the gene encoding galactosylceramidase (also known as “galactocerebrosidase” or “GALC”). Clinically, KD may present as a spectrum ranging from infantile-onset (IOKD) to later-onset forms (LOKD). People with IOKD usually present in the first months of life with rapid neurodegeneration and a fatal outcome, while people with LOKD may develop a more insidious leukodystrophy and peripheral neuropathy. Treatment for IOKD involves hematopoietic stem-cell transplantation (HSCT), which, when performed in the pre-symptomatic stage of the disease, has been shown to slow or prevent further central nervous system degeneration, although slower or more limited leukodystrophy as well as peripheral neuropathy still occur.

The pathophysiology of KD stems from the critical role of GALC for the catabolism of myelin glycolipids, by cleaving galactosylceramide (the simplest of galactosyl-sphingolipids that make up to one fourth of the dry weight of myelin ^1^) into free galactose and ceramide. Ceramide is then further processed by acid ceramidase into a free fatty acid and sphingosine. When galactosylceramide accumulates because of a deficient GALC activity, it is deacylated directly by acid ceramidase, leading to the formation of galactosylsphingosine (commonly known as “psychosine”)^2^. Psychosine exerts its cytotoxic effects through multiple mechanisms, including direct damage to cellular membranes due to its disruption of organellar organization^3,4^ and interaction with caspase and other pro-apoptotic proteins ^5–8^. Ultimately, accumulation of psychosine in oligodendrocytes leads to cell death and subsequent demyelination, accounting for the characteristic neurological manifestations of the disease.

Despite significant advances in the understanding of psychosine toxicity, the broader metabolic consequences of GALC deficiency remain poorly characterized. The advent of stable isotope metabolomics has revolutionized our ability to interrogate metabolic patterns and identify previously unrecognized metabolic perturbations in disease states. This approach enables precise tracking of nutrient utilization and metabolite synthesis, providing insight into the dynamic nature of cellular metabolism that cannot be captured by traditional metabolomic approaches.

In this study, we employed stable isotope small molecule metabolomics, lipidomics, and N-glycomics to systematically characterize metabolic patterns in a human cell-based KD model^9^. We uncover two key metabolic pathway changes associated with KD and nominate two potential therapeutic targets that could inform future treatment strategies for this devastating neurodegenerative disorder.

## RESULTS

Normally functioning GALC is responsible for removing galactose from galactosylceramides (Figure 1A). We first wanted to test the hypothesis that loss of GALC function would be accompanied by less glycolytic energy metabolism, since glucose-derived carbons could be used to compensate for decreased galactose recycling (Figure 1B). To examine this possibility, we used stable isotope metabolomics, where we supplied wildtype and *GALC* KO MO3.13 cells^9^ with ^13^C-labelled glucose. We used mass spectrometry to quantify metabolite pool size and utilization of carbon from ^13^C-glucose at typical cell culture concentration (25 mM, or “high Glc”) or closer to physiological concentration (2.5 mM, or “low Glc”). We observed that at physiologic (“low Glc”) glucose levels, *GALC* KO cells did exhibit differences in glycolytic metabolite levels, including a significant increase in the amounts of fructose 1,6-bisphosphate (F1,6BP) and 3-phosphoglycerate (3PG) (Figure 2A, Fig S1A, Table S1), but unchanged levels of ^13^C-glucose-derived labeling in these metabolite pools (Figure S1B, Table S2). *GALC* KO cells had lower ratios of high-energy carrying ATP (versus lower energy ADP and AMP) compared with control cells at high glucose, but not at low glucose (Figure S1C, Table S1). These observations indicate that there are alterations in glycolysis associated with *GALC* KO, however, decreased glycolysis may not be an important feature of the disease under physiological conditions. Instead, our observations suggest that metabolic pathways that depend on early glycolytic metabolites, such as glycosylation or serine biosynthesis, would have increased substrate availability and activity. It was indeed observed that metabolites associated with glycosylation accumulate at both high and low glucose concentrations for *GALC* KO cells (Figure 2B, Table S1). In the same conditions where these metabolite pool sizes increased, we observed increased utilization of carbon from ^13^C-glucose (Figure 2C, Table S2). Subsequent measurements to resolve levels of glycosylation-associated metabolite pools detected significantly less Gal1P in *GALC* KO cells grown in low glucose (Figures S1D). These observations are consistent with increased compensatory synthesis and/or decreased utilization of glycosylation metabolites.

**Figure 1.**
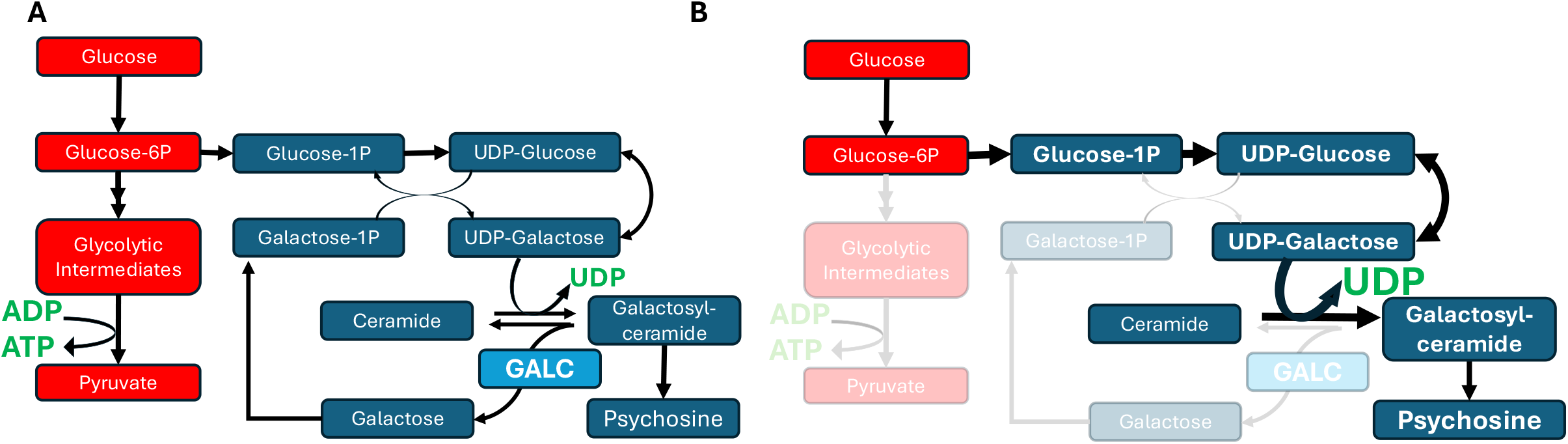
Schematic of psychosine synthesis in Krabbe disease. A) In healthy individuals, the GALC enzyme removes galactose from galactosylceramide. B) In individuals with Krabbe disease, the GALC enzyme is mutated, leading to accumulation of psychosine. We hypothesize that loss of GALC function would be accompanied by decreased free galactose, increased UDP levels, and more diversion of Glucose-6P into anabolic glycosylation-related pathways and away from catabolic glycolysis energy metabolism.

**Figure 2.**
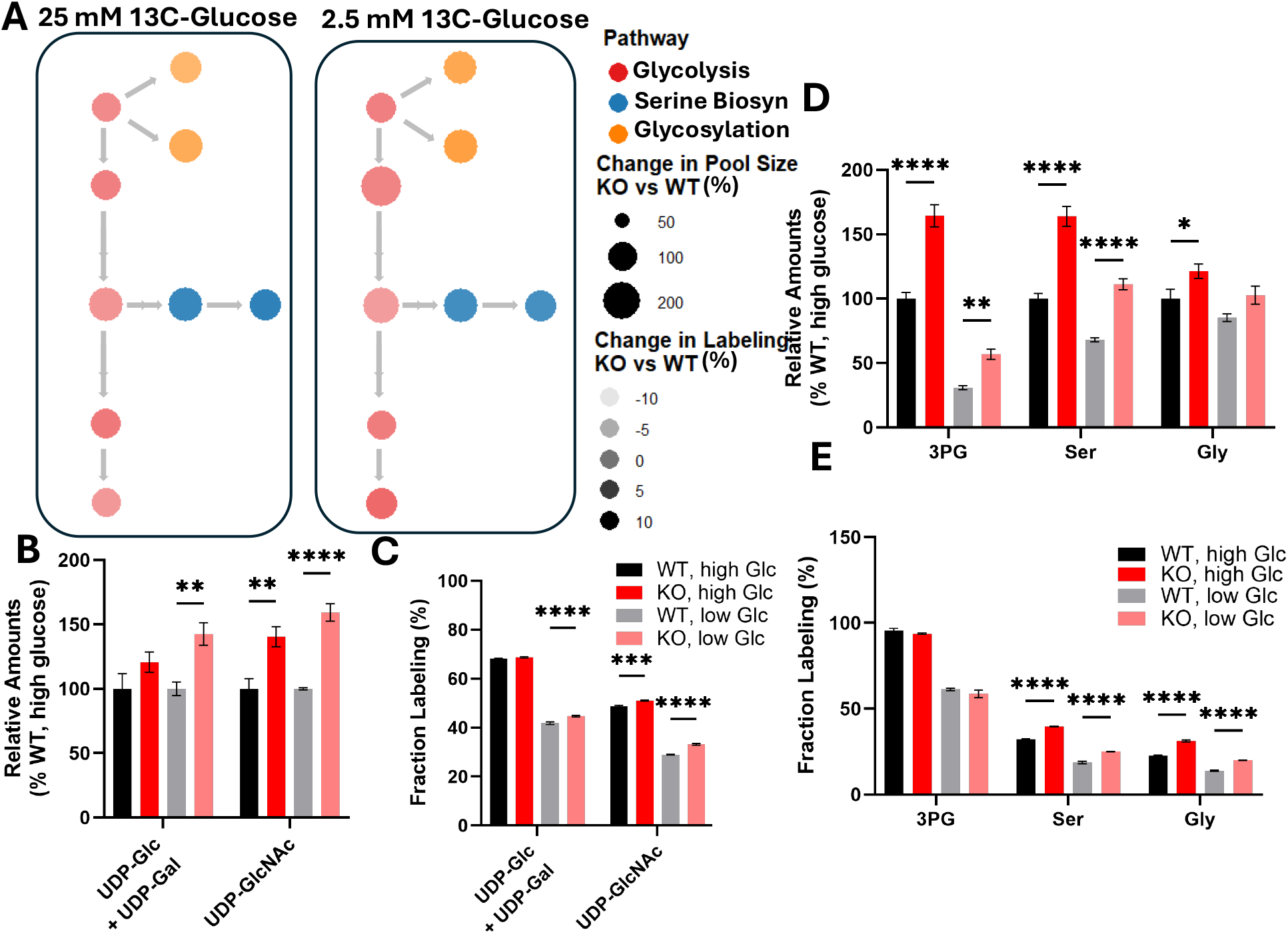
Stable isotope labeling reveals metabolic defects in a Krabbe disease model. A) Network plot depicting relative abundance and ^13^C-glucose-derived labeling of metabolite pools. Particularly at 2.5 mM glucose, there is an accumulation (increased radius) of early glycolytic metabolites (red) in *GALC* KO cells versus WT. Biosynthetic pathways including serine biosynthesis (blue) and glycosylation (orange) have elevated labeling derived from ^13^C-glucose (darker shading) in KO cells versus WT, at both glucose levels. Quantification of relative amounts (B) and ^13^C-glucose-derived labeling (C) in glycosylation-related metabolites reveals elevated levels and labeling with *GALC* KO, ****p<0.0001, ***p<0.001, **p<0.01 by 2-way ANOVA with Tukey’s correction for multiple comparisons. Quantification of relative amounts (D) and ^13^C-glucose-derived labeling (E) in serine biosynthesis-associated metabolites reveals elevated levels and labeling with *GALC* KO, ****p<0.0001, **p<0.01, *p<0.05 by 2-way ANOVA with Tukey’s correction for multiple comparisons. Data compiled from n = 4 replicates.

The glycolytic metabolite 3PG, which was increased in its concentration, but not in its ^13^C-labeling, in the *GALC* KO cells, is a precursor for serine biosynthesis. We also observed that serine and glycine metabolite pools increased in both size (Figure 2D, Table S1) and in proportion of ^13^C-glucose-derived labeling (Figure 2E, Table S2) at both glucose concentrations despite a lack of change in ^13^C -labeling of 3PG. This observation is consistent with increased activation of serine biosynthesis accompanying *GALC* KO.

Metabolomics also identified an accumulation of nucleotide sugars in *GALC* KO cells. To understand the downstream impact of this increase, we quantified abundance of a clinically relevant panel of glycans using time-of-flight mass spectrometry (Table S3). Individual glycans were categorized by shared structural characteristics and galactose composition. We observed that *GALC* KO was associated with an increase in biantennary glycans containing no galactose and a decrease in biantennary glycans with two galactose residues (Figure 3A, Table S4), consistent with glycan hypogalactosylation, as well as with increases in bisected glycans containing either one or two galactose residues (Figure 3B, Table S5), and no difference in monoantennary (Figure 3C, Table S5). The differences in these categories of glycans was driven by significant differences in several individual glycans (Figure 3D, Table S5) – most notably, by a decrease in Neu5Ac2Hex5HexNAc4 (m/z 1268), a mature biantennary glycan with two galactose residues, in *GALC* KO cells. In contrast, *GALC* KO was associated with an increase in Hex3HexNAc4 (m/z 814), an immature biantennary glycan with no galactose. This glycan serves as a precursor to galactosylation of biantennary glycans, thus increases in its abundance are consistent with a lack of availability of galactose (*i*.*e*., reflective of decrease site occupancy by galactose). As one of the most significantly increased glycans in our panel, Hex3HexNAc4 may represent a valuable biomarker for KD. Other decreased glycans included Fuc1HexNAc2 (m/z 442), Hex6HexNAc2 (m/z 855), and Hex7HexNAc2 (m/z 936), while other increased glycans included Hex4HexNAc3 (m/z 794), Hex9HexNAc2 (m/z 1098), Hex10HexNAc2 (m/z 1179), Hex11HexNAc2 (m/z 1260), along with bisected glycans Fuc1Hex4HexNAc7 (m/z 1274), Neu5Ac2Fuc1Hex5HexNAc5 (m/z 1443), Hex5HexNAc7 (m/z 1282), and Hex4Fuc1HexNAc5 (m/z 1070). Notably, elevated Hex3HexNAc4, as well as increases in other hypogalactosylated biantennary glycans, have been observed in known congenital disorders of glycosylation (CDG) treatable with galactose supplementation such as SLC35A2-CDG ^10,11^. To investigate whether galactose supplementation could ameliorate the glycosylation phenotype of the *GALC* KO cells, these were supplemented with either 1 mM or 5 mM galactose supplementation (as per the supplementation ranges studied in SLC35A2-CDG and PGM1-CDG^12,13^), with the 5 mM supplementation leading to a correction of the phenotype through decreased abundance of hypogalactosylated species (such as Hex3HexNAc4) and a corresponding increase in abundance of mature, galactosylated species (such as Neu5Ac2Fuc1Hex5HexNAc4) (Figure 3E, 3F, Table S6). Given that galactose is one of the precursors for galactosylceramide (and thus, for psychosine), we then measured psychosine concentrations in cells after galactose supplementation, and did not observe any increases (Figure S1E).

**Figure 3.**
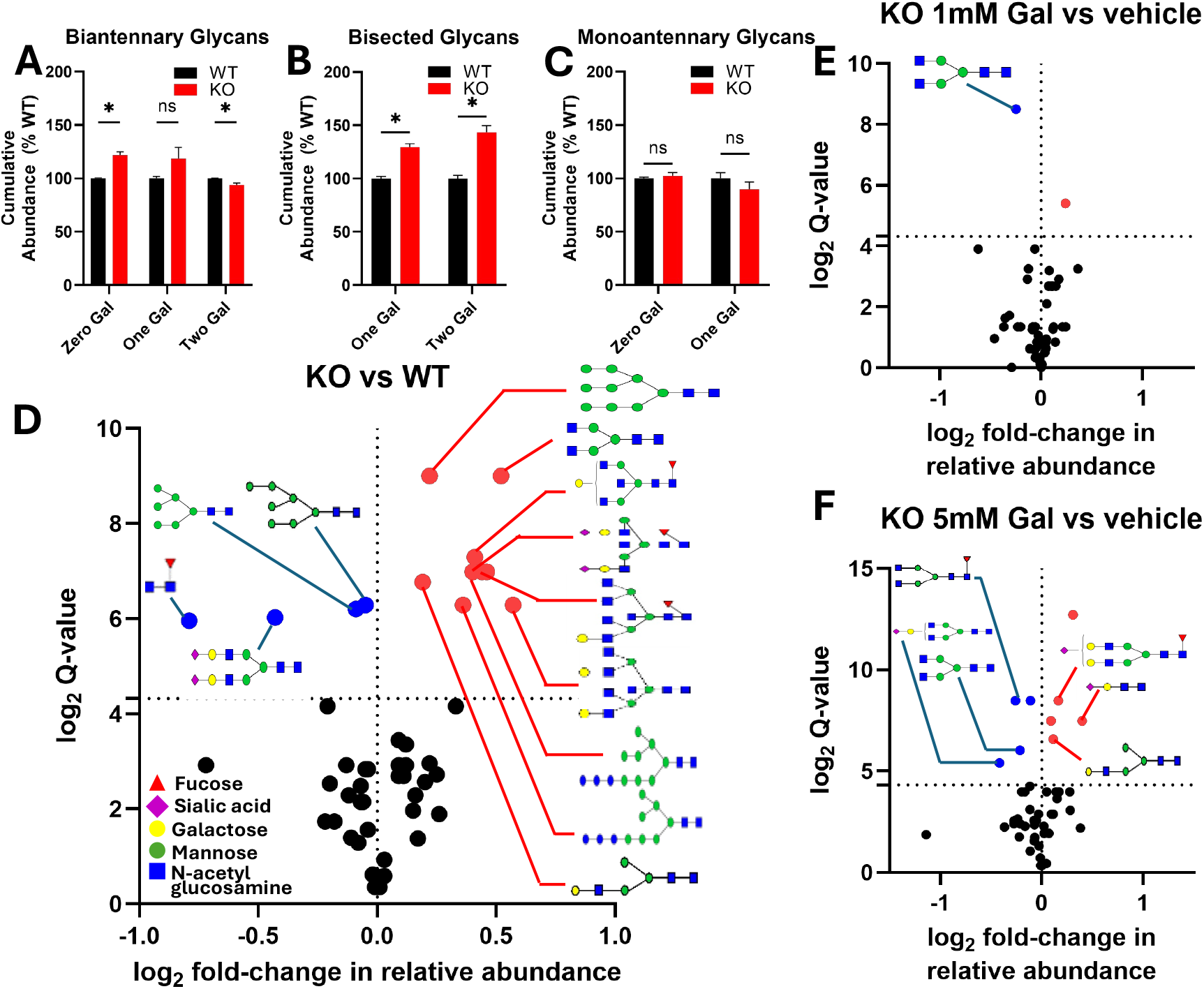
N-glycan profiling reveals hypogalactosylation associated with *GALC* KO. N-glycans abundance was normalized against their abundance in WT cells for a given experiment. N-glycans were grouped by structural characteristics, including those that are (A) biantennary, (B) bisected, or (C) monoantennary, and stratified by number of galactose. *GALC* KO cells had significantly more biantennary glycans with zero galactose, and significantly fewer biantennary glycans with two galactose. Bisected glycans included in our panel were elevated for *GALC* KO cells compared to controls, while monoantennary glycans did not differ. *p<0.05 by unpaired t-test. Data compiled from n=4 replicates. D) Volcano plot depicting N-glycans that differed between *GALC* KO cells and WT controls. Individual glycans were stratified by the log-fold abundance change relative to WT cells (x-axis) and the base 2 log-transformed q-value (y-axis, multiple t-tests corrected for multiple comparisons by two-stage step-up false discovery correction method of Benjamini, Krieger and Yekutieli). E-F) Volcano plots depicting N-glycans that shift in abundance relative to vehicle-treated KO cells with either 1mM or 5mM galactose supplementation. Data compiled from n=4 replicates.

Since serine is an important substrate in *de novo* ceramide synthesis, and that 3PG, serine, and glycine concentrations were increased in *GALC* KO cells, we hypothesized these changes may reflect an upregulation of the *de novo* ceramide synthesis pathway caused by a lack of galactosylceramide recycling (*i*.*e*., by “galactosylceramide trapping of galactose”). As tezacaftor has recently been identified as a well-tolerated inhibitor of ceramide synthesis through sphingolipid Δ^4^-desaturase (DEGS) ^14^, we tested whether this overactive metabolic pathway could be targeted therapeutically to decrease psychosine levels (Figure 4A). Tezacaftor was prioritized given that it is already FDA-approved for the treatment of cystic fibrosis. We quantified incorporation of carbons from ^13^C-glucose into ceramide pools using mass spectrometry. The peak area ratio corresponds to the amount of ^13^C-glucose containing ceramide relative to the total ceramide pool of that ceramide species. We observed that for some species, including ceramide (16:0), ceramide (18:0), and ceramide (24:1), which are direct precursors of galactosylceramide biosynthesis (Figure 4B-F), incorporation of carbons from glucose into ceramide pools increased over time and could be decreased with tezacaftor. Notably, the peak area ratios indicate that nearly two-thirds of the newly synthesized ceramide (24:1) pool incorporate carbons from glucose, suggesting that *de novo* ceramide synthesis is a major contributor to galactosylceramide (and subsequently psychosine) synthesis both in WT and *GALC* KO cells. Lastly, we observed that psychosine levels in *GALC* KO cells could be normalized by treatment with tezacaftor (Figure 4G).

**Figure 4.**
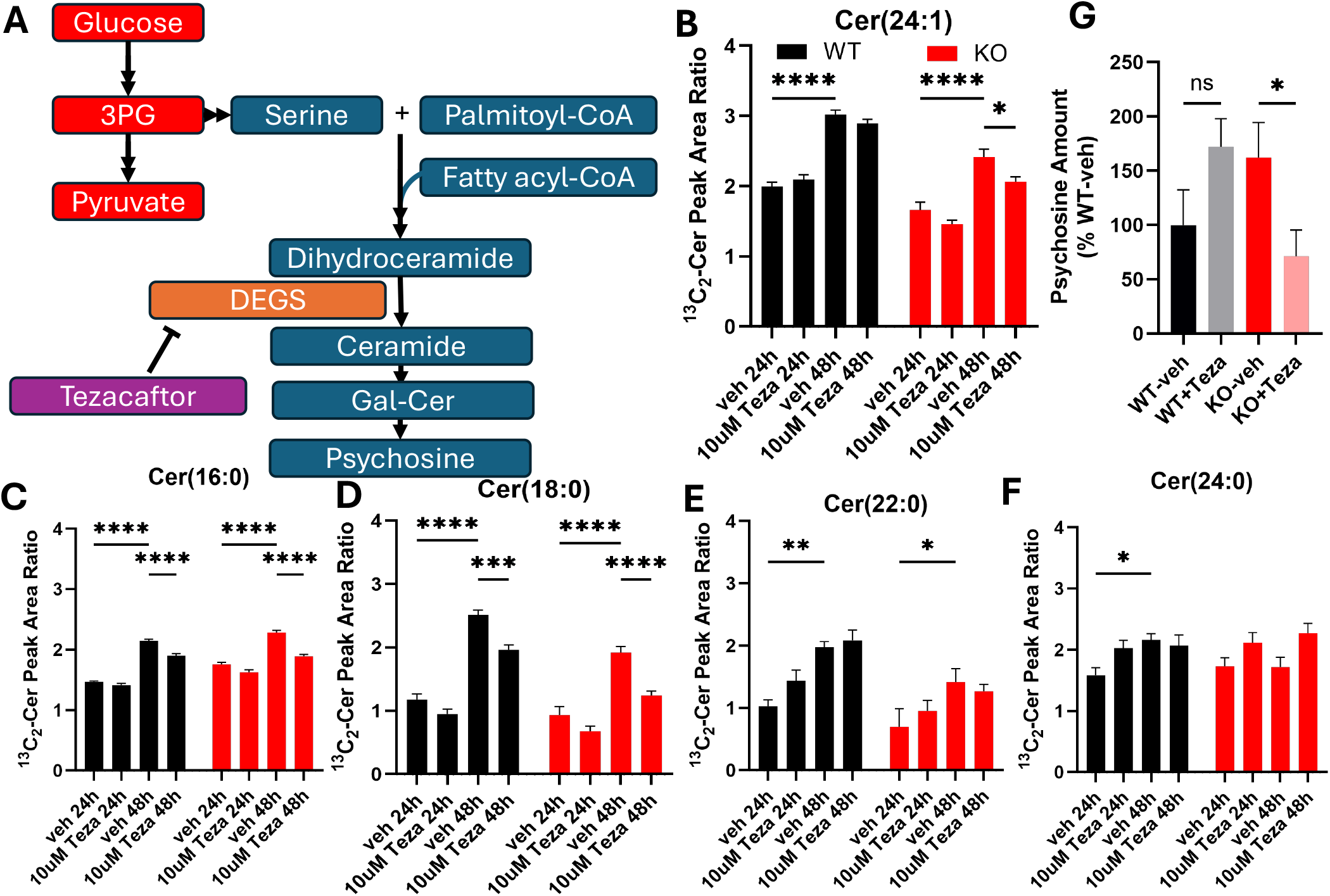
Tezacaftor inhibits de-novo synthesis of psychosine and psychosine precursors in *GALC* KO cells. A) Schematic of the target of Tezacaftor in the de novo biosynthetic pathway of ceramides. B-F) Peak area ratio refers to the ratio of ^13^C2-Cer from ^13^C6-glucose to natural ^13^C2-Cer. Quantification of contribution of ^13^C-glucose to Cer(24:1), a precursor to psychosine. Cer(24:1) labeling from ^13^C-glucose increased from 24h to 48h for both WT and *GALC* KO cells. Cer(24:1) labeling was significantly decreased in *GALC* KO cells treated with 10 µM Tezacaftor for 48h, compared with cells treated with vehicle control. Cer(16:0) and Cer(18:0) labeling was significantly decreased in *GALC* KO cells treated with 10 µM Tezacaftor for 48h, compared with cells treated with vehicle control. ****p<0.0001, ***p<0.001, **p<0.01, *p<0.05 by 2-way ANOVA with Tukey’s correction for multiple comparisons. Not significant comparisons not shown. Data compiled from n=6 replicates. G) Quantification of psychosine levels with Tezacaftor treatment. GALC KO cells have significantly lower levels of psychosine after 48h treatment with 10 µM Tezacaftor, compared with vehicle-treated cells. *p<0.05 by Student’s t-test. Data compiled from n=6 replicates.

## DISCUSSION

KD has classically been regarded as a “lysosomal *storage* disease”, with a focus on the build-up of undegraded material leading to cellular dysfunction through lysosomal engorgement and rupture. Although this understanding of KD has been substantiated by the “psychosine hypothesis” of toxicity, it may not account for the full pathophysiology. This approach has served as the basis for most therapeutic development in KD so far; nonetheless current therapeutic options for KD remain limited, with HSCT representing the primary treatment modality when performed early in the disease course. Despite being effective in extending lifespan in people with infantile-onset Krabbe disease (IOKD)^15^, and potentially in people with LOKD as well^16,17^, HSCT not only has a narrow window to be performed, but also may not completely halt the progression of disease - with ongoing developmental delays, spasticity, peripheral and cranial neuropathy reported in transplanted patients –reflecting incomplete resolution of the underlying pathology ^18–20^; in fact, a post-mortem analysis of a teenager with IOKD has shown evidence of continuing central nervous system white matter damage despite HSCT performed in the first month of life^21^. Long-term follow-up studies have shown that mortality is still higher than 20% at 10 years post-transplant ^22^. While experimental approaches including gene therapy and substrate reduction therapy (SRT) have shown promise in animal models, translation remains limited, and the need for novel therapeutic strategies remains urgent.

Recent metabolomic profiling studies in the *twitcher* (Twi) mouse model of KD have revealed complex metabolic perturbations extending beyond sphingolipid metabolism, including alterations in glucose energy metabolism^23,24^. Similarly, recent experiments on *galcb* KO zebrafish (which mimic the pathology of KD) have shown a potential psychosine-independent toxic mechanism of lactosylceramide accumulation^25^. These findings suggest that the pathogenic mechanisms underlying KD may be more complex than previously appreciated, involving other complex molecules, and potentially small-molecule metabolism. Our previous work highlights glycolysis as a central hub between anabolic and catabolic cell processes^26^ – here, we used stable isotope labeling approaches to investigate how perturbed glucose energy metabolism might impact glycolysis-associated biosynthetic pathways.

Our findings also suggest that defective glycosylation may play a role in the pathophysiology of KD, as there was significant hypogalactosylation of N-glycans in the KO cells. The pattern of hypogalactosylation was similar to the one observed in congenital disorders of glycosylation (CDGs) that impact UDP-galactose availability in the Golgi apparatus, such as SLC35A2-CDG, with an increase of truncated bi-antennary glycans that terminate one or both antennae in GlcNAc^10,27,28^. As antennary GlcNAc (ie., GlcNAc residues that are distal to the trimannose N-glycan core) are necessarily followed by Gal residues, which then should be followed by terminal sialylation, GlcNAc-terminated glycans likely reflect decreased availability of UDP-Gal as a galactosyl donor. In the context of our findings on metabolic analysis of a disruption of early glycolysis as well as increased serine biosynthesis and incorporation into *de novo* sphingolipids, this likely reflects decreased recycling of Gal from GalCer and psychosine which is compensated by a shunting of UDP-Gal to sphingolipid synthesis instead of glycosylation. This points to a similar pathophysiological process between SLC35A2-CDG and KD, reflecting the tight connection between galactose and myelin glycosphingolipid metabolism^13,29^. The accumulation of high-mannose glycans (Hex9HexNAc2, Hex10HexNAc2, and Hex11HexNAc2) may be a consequence of Hex3HexNAc4 accumulation. Elevated Hex3HexNAc4 coupled with blocked biantennary glycan synthesis may then result in elevated levels of the bisected glycans observed. Although it remains to be proven whether the glycosylation abnormalities drive some of the pathology of KD (*i*.*e*., whether KD is also a CDG), this leads to the hypothesis that galactose supplementation, which has been shown to lead to clinical and biochemical improvements in people with SLC35A2-CDG^30^ (as well as the biochemically adjacent CDG PGM1-CDG^12,31^), may lead to improvements in people with KD. Given that oligodendrocyte-specific hypogalactosylation has been shown to lead to hypomyelination, seizures, and gliosis in a model of SLC35A2-CDG^32^, it is possible that correcting the hypogalactosylation – which we have shown feasible, at least *in vitro*, by galactose supplementation – could lead to clinical improvements in patients with KD, although further studies in model organisms are necessary before translating this idea into clinical studies. A safety aspect of this approach was to ascertain whether galactose supplementation could lead to an increase in production of psychosine, as UDP-Gal is necessary for GalCer synthesis; specially as a recent study showed that Gal supplementation in fibroblasts of people with SLC35A2-CDG led to increased glycosphingolipid production^13^. In our experiments, we have shown that galactose supplementation did not increase psychosine concentration, which increases its promise as a potential therapeutic.

Further exploration of our metabolomic data led us to identify an increased *de novo* synthetic flux of ceramides, likely also caused by “trapping” of ceramide as GalCer. This highlights the potential therapeutic role of inhibitors of ceramide synthesis in KD, an approach known as “substrate reduction therapy” (SRT)^33^. SRT has been tested for KD in the past: L-cycloserine, a serine-palmitoyltransferase (SPT) inhibitor, has been shown to extend survival and attenuate neuropathology in Twi mice^34,35^. A different compound, BMN-S202, has shown promise by inhibiting ceramide galactosyltransferase^9,36^. Both these compounds, however, have been limited to cell and animal studies due to the toxicity and incomplete correction of pathology. In this study, we used a new approach to SRT in KD by using a well-tolerated, FDA-approved drug – tezacaftor – that inhibits DEGS, and showed that it normalizes psychosine levels in treated *GALC* KO cells. Using an FDA-approved treatment with known safety and dosing may allow for easier translation into clinical studies if efficacy is demonstrated in animal models.

A limitation of our study includes the use of an immortalized cell line that includes a rhabdomyosarcoma fusion may alter the underlying biochemistry of the base oligodendrocytes used. The lack of a pluricellular population may also limit how generalizable our findings are, given that *in vivo* oligodendrocytes interact biochemically with astrocytes, microglia, and neurons. to address these concerns, experiments for replications of our findings are underway on a mouse model of the disease, as well as on banked brain species of patients with KD.

## CONCLUSION

The metabolic abnormalities underlying the pathogenesis of KD are more complex than simple psychosine accumulation. In this paper, we have used multi-omics including lipidomics, N-glycomics, and labeled small-molecule metabolomics to investigate the role of galactose and serine trapping due to GALC deficiency in a validated human oligodendrocyte-based model of KD. There was increased flux through the serine biosynthesis pathway leading to shunting of 3-phosphoglycerate into *de novo* sphingolipid synthesis; this was attenuated by tezacaftor administration, which also normalized psychosine levels. We have also shown that there is an increased use of exogenous Glc-derived UDP-Gal as well as global N-linked hypogalactosylation, likely reflecting a decrease in Gal recycling from GalCer and psychosine leading to decreased UDP-Gal availability; and that this was corrected by Gal supplementation with no corresponding increase in psychosine. Overall, our findings open new avenues for the understanding and treatment of KD.

## Methods

### Cell Culture

MO3.13 *GALC* WT and KO cell lines have been previously validated^9^. Cells were cultured in DMEM (Gibco) supplemented with 20% FBS (Corning) and 1% Penicillin-streptomycin. Cells were allowed to grow to ∼80% confluence before subculturing using 0.25% Trypsin-EDTA (Thermo-Fischer). Cells were cultured under standard incubator conditions. Prior to metabolomic, lipidomic, or N-glycomic analyses, cells were plated at a density of 200,000 cells/cm^2^ for 24h before exchanging media for complete media containing 20% heat-inactivated FBS. FBS was heat-inactivated at 60 °C for 30 minutes, followed by immediate cooling in an ice bath. For experiments with controlled glucose or stable-isotope labeled glucose, heat-inactivated media was made using glucose- and pyruvate-free DMEM (Gibco), supplemented with D-(+)-glucose (Sigma) or D-Glucose (U-^13^C_6_, 99%, Cambridge Isotope Laboratories) and pyruvate (Fisher Scientific).

### Polar Metabolite Extraction and UHPLC-MS analysis

Cell pellets containing 500,000 cells were washed with ice-cold PBS, followed by a 60-minute incubation with 80% methanol (Sigma) at -80 °C. After centrifugation, supernatant containing extracted polar metabolites was dried using a speedvac. Dried metabolites were resuspended in 100 ul 50% ACN:water and 5 µL were loaded onto a Luna 3um NH2 100A (150 × 2.0 mm) column (Phenomenex). The chromatographic separation was performed on a Vanquish Flex (Thermo Scientific) with mobile phases A (5 mM NH4AcO, pH 9.9) and B (95% ACN, 5% H2O) and a flow rate of 200 μL/min. A linear gradient from 10% A to 95% A over 18 min was followed by 7 min isocratic flow at 95% A and reequilibration to 10% A. Metabolites were detected with a Thermo Scientific Q Exactive mass spectrometer run with polarity switching in full scan mode with an m/z range of 70-975 and 70.000 resolution. Maven (v 8.1.27.11) was utilized to quantify the targeted metabolites by AreaTop using accurate mass measurements (< 5 ppm) and expected retention time as previously verified with standards.

C13 natural abundance corrections were made using AccuCor. Relative amounts of metabolites were calculated by summing up the intensities of all detected isotopologues of a given metabolite. Data analysis was performed using in-house R scripts.

For analysis using an Ion Chromatography System (ICS) 5000 (Thermo Scientific), the extract was diluted 5-fold with 50% ACN and 10 ul was loaded onto a Dionex IonPac AS11-HC-4 μm anion-exchange column using a flow rate of 350 μL/min. KOH concentrations in LC method: 0 min, 5 mM; 8 min, 35 mM; 13 min, 95 mM; 18 min, 95 mM; 18.1min, 5 mM; 23min, 5 mM. Post column, MeOH was infused at 60 ul/min. Using a Q Exactive mass spectrometer (Thermo Scientific), full SIM data was acquired at 70K resolution in negative polarity mode with a scan range of 70-900 m/z. Data extraction was conducted using Maven software.

### Measurement of N-glycan profiles in cell samples

N-glycan preparation was carried out with a RapiFluor-MS™ N-Glycan Kit following the manufacturer’s instructions. Briefly, 11 µL of cell lysate (sonicated in water), 3 µL of [^13^C] sialylylglycopeptide (100 µg/mL), 6 µL of buffered RapiGest SF™ solution (5%, w/v), were mixed, heated at 90°C for 10 min and cooled to room temperature (RT). 1.2 µL of Rapid PNGase F™ was added to the mixture and incubated at 52°C for 5 min and cooled to RT. 12 µL of RapiFluor-MS™ Reagent in anhydrous dimethylformamide (6.9%, w/v) was added, incubated at RT for 5 min, and then diluted with 358 µL of acetonitrile. A HILIC 96 well µElution™ plate operated on a vacuum manifold was used to isolate N-glycans. The wells were conditioned with 200 µL of water, then equilibrated with 200 µL of 85% acetonitrile/water (v/v). The sample was loaded into the wells and washed twice with 600 µL of 1% formic acid in 90% acetonitrile/water (v/v), before elution with 3x 30 µL of elution buffer and subject to QTOF analysis as described previously^37^.

### Measurement of ceramides and ^13^C_2_-ceramides, and psychosine in cell samples

Cells (1 million) were suspended in 0.25 mL PBS. Ceramides were extracted from 50 µL of the cell suspension using the Bligh–Dyer extraction method^38^. Quantification of ceramides and ^13^C_2_-ceramides was performed using a Shimadzu Prominence HPLC system (Columbia, MD) coupled to an Exploris 120 mass spectrometer (Thermo Fisher Scientific, Waltham, MA). LC separation was achieved on a Halo C8 column (4.6 mm × 100 mm, 2.7 µm) at a flow rate of 0.8 mL/min. Mobile phases consisted of (A) 5 mM ammonium acetate in water and (B) 5 mM ammonium acetate in isopropanol–methanol (1:1, v/v). The gradient program was as follows: 0–40 min, 60– 95% B; 40–55 min, 95% B; 55–55.1 min, 95–60% B; and 55.1–65 min, 60% B. The column temperature was maintained at 50 °C, and the injection volume was 5 µL.

The mass spectrometer was equipped with a heated electrospray ionization (HESI) source and operated at a resolution of 120,000, with sheath gas at 60, auxiliary gas at 20, sweep gas at 5, ion transfer tube temperature at 275 °C, vaporizer temperature at 450 °C, and spray voltage of 5,000 V. Data were acquired over an m/z range of 500–1,000 with an AGC target of 1 × 10^6 and automatic maximum injection time. Peak areas for ceramides and ^13^C_2_-ceramides were integrated using Xcalibur (version 4.6.0.100). Natural ^13^C_2_-ceramide contributions were calculated from ceramide peak areas based on theoretical isotopic abundances and subtracted from measured ^13^C_2_-ceramide signals. Final results were reported as the peak area ratio of ^13^C_2_-ceramides derived from ^13^C_6_-glucose to total ceramides. Psychosine in cell samples was measured by LC– MS/MS as previously described.^39^

**Supplementary Figure 1.**
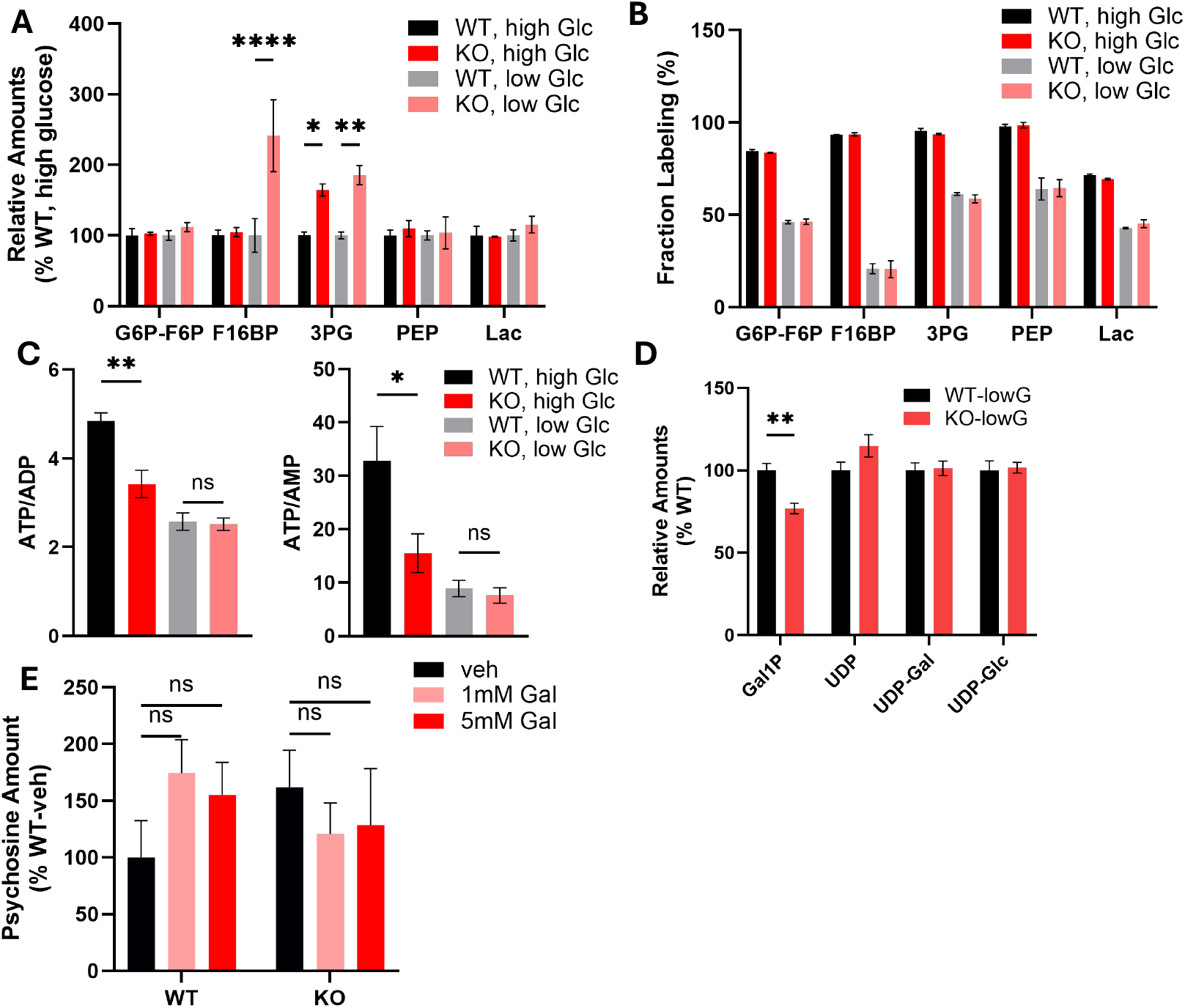
Glycolysis, glycosylation, and energy balance are perturbed in *GALC* KO cells. Glycolytic metabolite pools’ (A) Relative amounts, normalized against the average level of WT cells treated with high glucose, and (B) ^13^C-glucose-derived labeling. These are the same metabolites, in order, presented in Figure 2A: Glucose-6-phosphate or Fructose-6-phosphate (G6P-F6P), Fructose1,6-bisphosphate (F16BP), 3-phosphoglycerate (3PG), Phosphoenolpyruvate (PEP), and Lactate (Lac). ****p<0.0001, **p<0.01, *p<0.05 by 2-way ANOVA with Tukey’s correction for multiple comparisons. Data compiled from n=4 replicates. C) Ratios of the pool sizes of energy-carrying species. The ratio of ATP to ADP and ATP to AMP was significantly lower in *GALC* KO cells compared to WT cells, but only at high glucose levels. **p<0.01, *p<0.05 by one-way ANOVA with Šidák’s correction for multiple comparisons. Data compiled from n=4 replicates. D) Glycosylation metabolite pools’ relative amounts, normalized against the average level of WT cells treated with low glucose. Gal1P levels were significantly reduced in *GALC* KO cells. **p<0.01 by 2-way ANOVA with Tukey’s correction for multiple comparisons. Data compiled from n=4 replicates. E) Psychosine levels in WT and KO cells do not increase with 1mM nor 5mM galactose supplementation. n.s. not significant by 2-way ANOVA with Tukey’s correction for multiple comparisons. Data compiled from n=6 replicates.

